# Microtubule Engagement by the WD40-Containing Tail of Kinesin-4 KIF21B

**DOI:** 10.64898/2026.07.29.741364

**Authors:** Aryan Taheri, Benjamin LaFrance, Julia Peukes, Ankit Rai, Anna Akhmanova, Eva Nogales

## Abstract

Kinesin tails are structurally diverse and mediate a range of functions, including autoinhibition, cargo binding, and microtubule regulation. Within the kinesin-4 family, KIF21A and KIF21B have emerged as key regulators of microtubule network organization and dynamics in neurons and immune cells, yet the molecular basis of this activity has remained unclear. Here, we combined single-particle cryo-electron microscopy (cryo-EM) and cryo-electron tomography (cryo-ET) to examine how the KIF21B tail engages microtubules. We find that conserved residues in the WD40 β-propeller and an adjacent N-terminal linker contact successive tubulin dimers along a single protofilament, forming an extended longitudinal binding mode that spans both the intradimer and interdimer interfaces. Cryo-EM 3D classification further revealed two distinct engagement states, a tilted state, in which the β-propeller makes partial contacts with the microtubule while the linker remains anchored, and a flat state, in which the β-propeller lies flush with the lattice surface. Cryo-ET of full-length KIF21B reveals multiple binding configurations on the microtubule lattice, including orientations consistent with crosslinking adjacent microtubules. Together, these findings provide a structural framework for tail-mediated microtubule attachment by a neuronal kinesin-4 and suggest how its distinctive WD40 domain may contribute to microtubule regulation and organization of microtubule arrays.

---

Kinesins are a superfamily of microtubule (MT) motors that drive diverse cellular processes, including long-range intracellular cargo transport, mitotic spindle organization, and regulation of MT dynamics (Vale et al., 1985; Yildiz, 2025). Each kinesin contains a conserved motor domain that couples ATP hydrolysis to MT binding and force generation. Based on sequence homology within this motor domain, kinesins are classified into 14 subfamilies (Wickstead and Gull, 2006). Much of the functional diversity among kinesins, however, arises from their tail regions, which vary extensively across families and determine cargo specificity, regulatory mechanisms, and the mode of MT interaction (Coy et al., 1999; Nithianantham et al., 2023).

Members of the kinesin-4 family are best known for their role in inhibiting MT growth during mitosis, ciliogenesis, and other processes across diverse cell types, including immune cells and neurons (Bringmann et al., 2004; Bieling et al., 2010; Subramanian et al., 2013; Yue et al., 2018; Hooikaas et al., 2020). All kinesin-4 members share a conserved N-terminal motor domain that drives motility and regulates MT dynamics, although two members, KIF7 and KIF27, are immotile or exhibit only limited motility (Yue et al., 2018). The KIF21B motor domain alone is sufficient to slow MT growth, and structural studies of the C. elegans ortholog bound to tubulin suggest that kinesin-4 motors promote a depolymerizing MT architecture by increasing tubulin dimer curvature at MT plus ends (Taguchi et al., 2022).

Within the kinesin-4 family, the KIF21 subfamily is distinguished by its tissue distribution and disease relevance. Human KIF21A is ubiquitously expressed, while KIF21B is enriched in the brain, spleen, and testes (Marszalek et al., 1999). Dominant gain-of-function mutations in KIF21A cause congenital fibrosis of the extraocular muscles types 1 and 3 (CFEOM1 and CFEOM3), neurodevelopmental disorders characterized by defective axon guidance (van der Vaart et al., 2013; Cheng et al., 2014; Al-Haddad et al., 2021). KIF21B knockout mice are viable but display abnormal neuron morphology, synaptic defects, and impaired learning and memory, and knockout neurons also show decreased MT growth rates and increased MT lengths (Muhia et al., 2016; Ghiretti et al., 2016). Elevated KIF21B expression has been associated with accelerated neurodegeneration in Alzheimer”s disease (Kreft et al., 2014), and gain-of-function missense variants in KIF21B have been identified in patients with agenesis of the corpus callosum and microcephaly, with expression of these variants inducing abnormal neuronal migration (Asselin et al., 2020). Together, these findings implicate KIF21B as a regulator of MT dynamics whose altered activity can disrupt neuronal migration, axonal organization, and brain morphogenesis.

Unlike their conserved motor domains, the tail regions of kinesin-4 family members are highly diverse, enabling distinct mechanisms of motor regulation, autoinhibition, and cellular specialization. The KIF4 tail binds chromosomes to support its function as a chromokinesin during mitosis (Mazumdar et al., 2004), whereas the KIF27 tail serves as a scaffold for ciliogenesis (Park et al., 2025). KIF7 and KIF21A/B harbor inhibitory elements in their tails that mediate autoinhibition through a folded-back conformation (Bianchi et al., 2016; Haque et al., 2026). KIF7 is autoinhibited in the cytosol and activated by Hedgehog signaling, whereas KIF21A/B autoinhibition is thought to be relieved by cargo binding (Weng et al., 2018; Haque et al., 2026). Among kinesin-4 motors, KIF21A and KIF21B are further distinguished by a WD40 β-propeller domain at the distal tail (Guo et al., 2018). In KIF21B, this domain promotes persistent association with MT plus ends by preferentially recognizing the GTP-tubulin state of the MT lattice, enhancing KIF21B-mediated MT growth pausing (van Riel et al., 2017). Although both the WD40 β-propeller and an adjacent N-terminal linker have been shown to contribute to MT binding, the molecular basis of this interaction has remained unknown.

Here, we define the structural basis of MT engagement by the KIF21B tail. Single-particle cryo-EM reveals that the WD40 β-propeller and its N-terminal linker together bind across successive tubulin dimers along a single protofilament, with the linker engaging the intradimer interface and the β-propeller bridging the interdimer interface. 3D classification identified two distinct engagement states: a tilted state in which the β-propeller makes partial contacts with the MT while the linker remains anchored, and a flat state in a more extensive engagement with the lattice surface. Cryo-ET of full-length KIF21B bound to MTs reveals binding configurations consistent with both single MT engagement and crosslinking of adjacent MTs. Together, our findings provide a mechanistic framework for understanding how the KIF21B tail may contribute to MT dynamics regulation and the organization of MT networks.

## RESULTS

### The distal KIF21B tail binds GMPCPP-stabilized MTs

To define how the KIF21B tail engages MTs, we purified a C-terminal tail fragment previously shown to bind MTs strongly and to preferentially decorate GMPCPP stabilized MTs (van Riel et al., 2017). This construct, termed KIF21BL–WD40, spans residues 1247–1674 and comprises an N-terminal linker followed by a WD40 β-propeller (Figure 1A,B), both of which were previously shown to contribute to MT binding (van Riel et al., 2017). Mass photometry of purified KIF21BL–WD40–mNeonGreen (mNG), expressed in Sf9 cells, confirmed that this fragment is a monomer in solution, with an expected mass of 71 kDa and a measured mass of 74 ± 10 kDa (Figure 1C). In MT decoration assays, KIF21BL–WD40–mNG bound GMPCPP stabilized MTs with an apparent dissociation constant of 180 ± 25 nM (Figure 1D).

**Figure 1.**
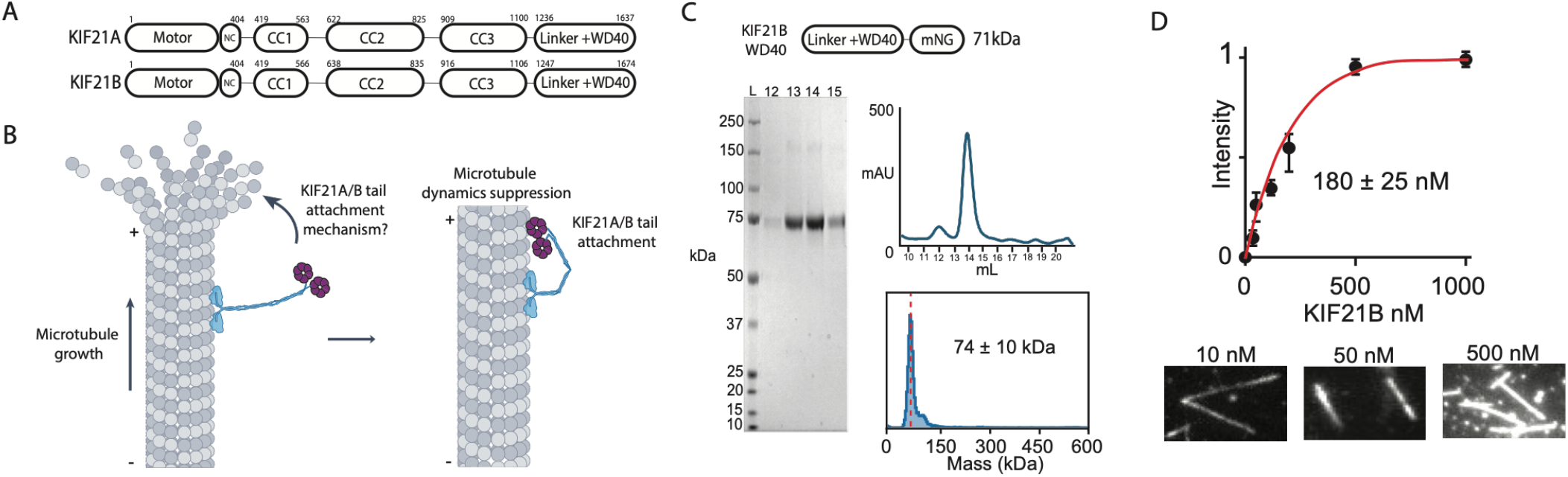
The distal KIF21B tail binds GMPCPP-stabilized MTs. (A) Domain maps of KIF21A and KIF21B highlighting major domains and residue boundaries. (B) Previously proposed model for MT dynamics regulation by KIF21B through attachment of the C-terminal tail to the MT tip. (C) (Top) Domain map of the KIF21BL–WD40–mNG construct. (Middle right) Size-exclusion chromatography profile of purified KIF21BL–WD40–mNG. (Left) SDS–PAGE analysis of size-exclusion fractions. (Bottom right) Mass photometry profile. (D) Affinity curve generated by quantifying fluorescence intensity on MTs as a function of protein concentration. (Bottom) Fluorescence images of KIF21BL–WD40– mNG bound to MTs at increasing concentrations.

### The KIF21B tail binds along protofilaments

Cryo-EM analysis of GMPCPP stabilized MTs incubated with KIF21BL–WD40–mNG revealed large, regularly spaced protein densities on the MT surface in 2D class averages (Supplementary Figure 1). 3D reconstruction and refinement yielded a map at an overall resolution of 3.1 Å. The structure shows that the WD40 β-propeller sits at the αβ-tubulin interdimer interface (Figure 2A,B), while a second density at the intradimer interface corresponds to a short, highly conserved segment immediately N-terminal to the β-propeller (Figure 2B,C). This arrangement provides a molecular explanation for previous deletion experiments showing that both the WD40 β-propeller and the adjacent linker region are required for robust MT binding (van Riel et al., 2017).

**Figure 2.**
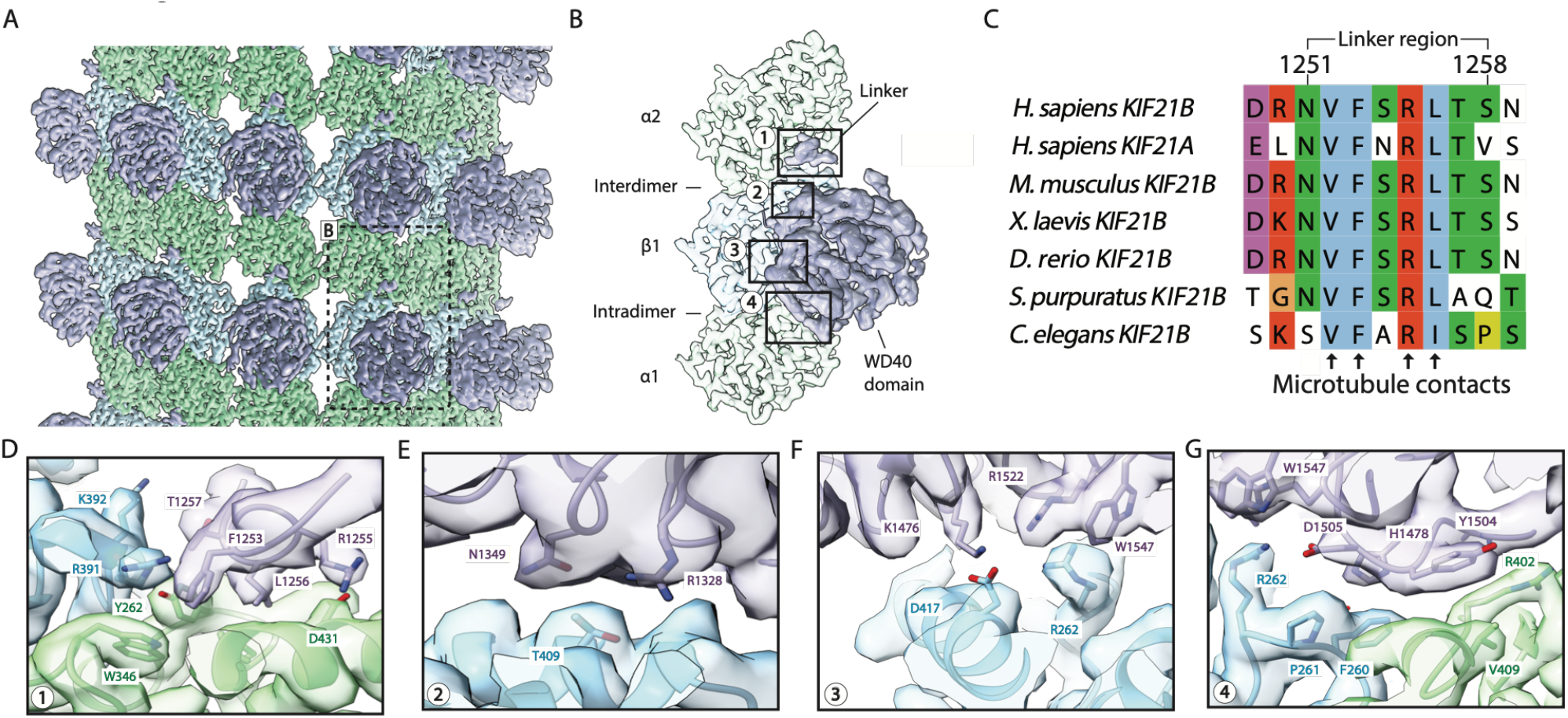
The KIF21B tail binds along protofilaments. (A) Cryo-EM reconstruction of MT bound KIF21BL–WD40– mNG. α-tubulin, β-tubulin, and KIF21B are shown in green, light blue, and light purple, respectively. (B) Density map obtained by symmetry expansion, highlighting the linker region and residues at the boundaries of the N-terminal linker and WD40 β-propeller. (C) Sequence conservation of the linker MT-binding region across species and with KIF21A. (D) Details of KIF21B linker engagement with the MT. (E–G) Details of KIF21B WD40 domain engagement with the MT.

### The KIF21B WD40 β-propeller bridges the MT longitudinal interface

The local resolution of the β-propeller density (~3–4.5 Å; Supplementary Figure 2) allowed us to build an atomic model and define its MT-binding footprint spanning both α- and β-tubulin (Figure 2D–G). On the face of the β-propeller proximal to the N-terminal linker, contacts are fewest and density is weakest (Figure 2G). KIF21B residues D1505, H1478, and Y1504 insert into the tubulin interdimer interface, contacting β-tubulin R262 and F260 and α-tubulin R402, respectively (Figure 2G). At the periphery, W1547 contacts β-tubulin R262, while R1522 and K1476 engage β-tubulin D427 (Figure 2F). Together, these residues form an extensive binding surface conserved between KIF21A and KIF21B, suggesting a shared tail–MT attachment mechanism within the KIF21 subfamily (Supplementary Figure 3). The numerous contacts spanning multiple tubulin dimers and the interdimer interface are consistent with the preference of this domain for the expanded, GTP-state MT lattice (van Riel et al., 2017; Alushin et al., 2014).

### Distinct engagement states of the KIF21B WD40 β-propeller

3D classification of the cryo-EM data identified two distinct binding states of the WD40 β-propeller. In the fully engaged “flat” state described above, the β-propeller lies flush with the MT surface and makes extensive contacts across the interdimer interface. In a second, “tilted” state, the β-propeller is rotated by ~35° relative to the flat state, with weaker local resolution consistent with greater conformational flexibility (Figure 3A–C). The largest displacement between states occurs at the α-tubulin and β-tubulin H12 contact, where the β-propeller shifts by a ~9Å rotation and a ~15 Å vertical displacement (Figure 3A,B). In the tilted state, the β-propeller forms a contact site on α-tubulin, with Y1504 and D1505 approaching E414 and R402 respectively. A distinct electrostatic interaction network forms between the KIF21B tail and β-tubulin H12, involving contacts between R1565 and R1522 with β-tubulin residues E421, Q424, and Y425 (Figure 3C).

**Figure 3.**
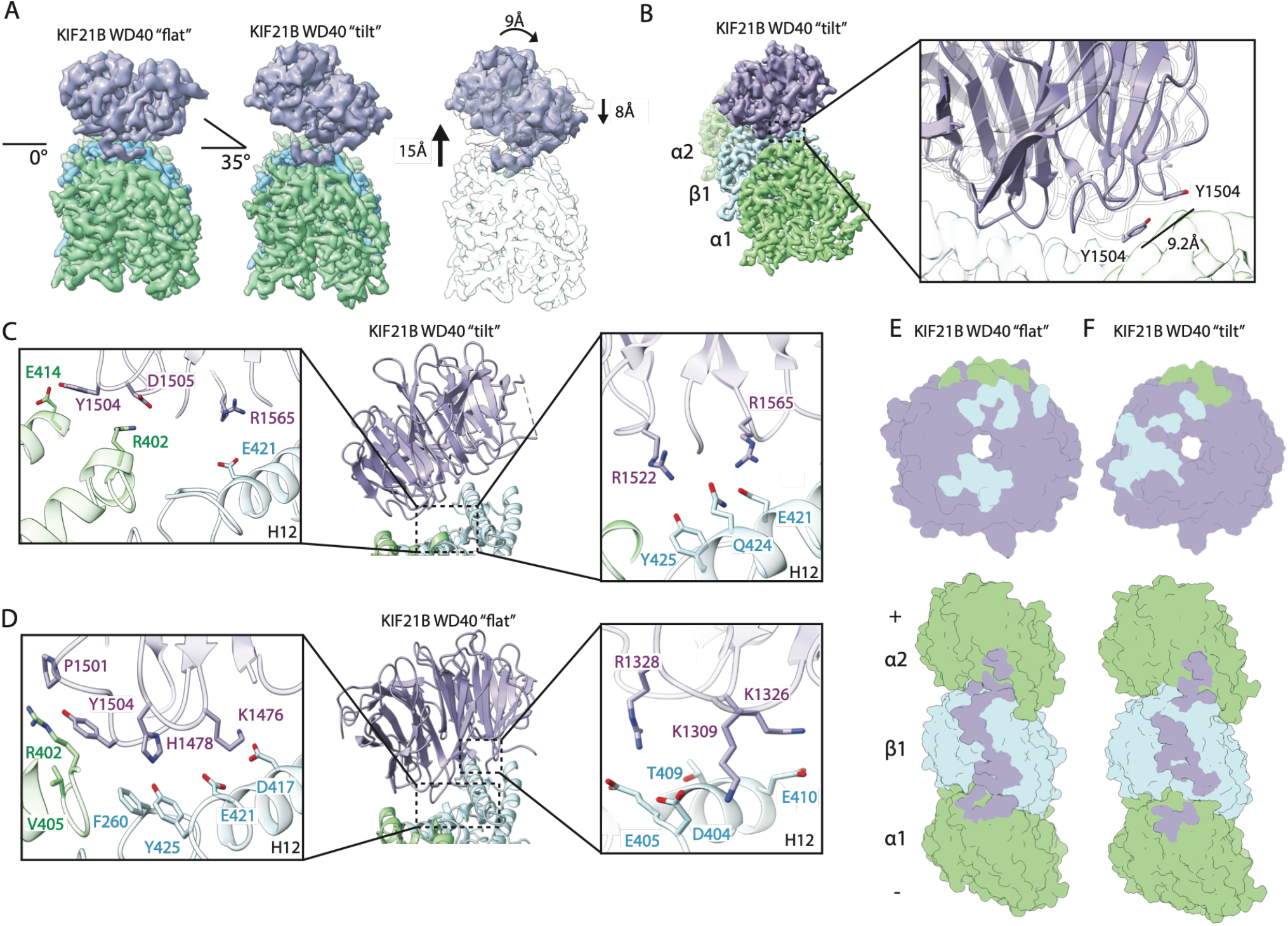
Distinct engagement states of the KIF21B WD40 β-propeller. (A) (Left) End-on view down a protofilament showing the flat conformation of the KIF21B β-propeller. (Middle) The tilted conformation of the KIF21B β-propeller on MTs. (Right) Overlay of tilted and flat conformations showing displacement measurements. (B) (Left) Tilted KIF21B WD40 density at the inter-protofilament interface, contacting β_3_ and α_2_-tubulin. (Right) Distance between Y1504 residues in the two conformations. (C,D) Close-up views of KIF21B–tubulin interfaces in the tilted (C) and flat (D) conformations. (E,F) Surface footprints of the KIF21B β-propeller on the MT in the flat (E) and tilted (F) conformations. (Top) β-tubulin and α-tubulin contacts on the β-propeller. (Bottom) KIF21B β-propeller contacts on the MT surface.

Notably, Y1504 adopts an alternative position at the interdimer docking site in the tilted state, approaching E414 in a configuration that disrupts neighboring contacts present in the flat state where it approaches R402 (Figure 3B,C,D). In contrast, the linker density at the intradimer pocket is similarly positioned in both states, despite the large displacement of the β-propeller. In the flat state, a larger interaction network is present in comparison to the tilted state, and the opposite face of the β-propeller contacts β-tubulin (Figure 3D,E,F). These observations are consistent with a model in which the linker remains anchored on the MT while the β-propeller samples tilted and flat configurations.

### MT binding configurations of full-length KIF21B

Full-length KIF21B contains two MT-binding modules, the motor domain and the WD40–linker region, and can therefore engage one or more MTs simultaneously. To visualize how full-length KIF21B interacts with MTs, we incubated purified full-length KIF21B with GMPCPP-stabilized MTs and analyzed the sample by cryo-ET. MTs exhibited 14 protofilaments consistent with GMPCPP polymerization and frequently appeared in bundles (Figure 4A,B). Within bundled regions, we observed protein densities associated with MT surfaces in multiple configurations, including densities positioned between adjacent MTs and densities bound along single MTs (Figure 4C–F). Bridging densities connecting neighboring MTs were frequently oriented at an angle rather than along the shortest orthogonal path, yet maintained an approximately constant length of 28 ± 2 nm despite variation in MT–MT spacing, consistent with a flexible crosslinking configuration (Figure 4D). Among particles bound to a single MT, we observed two sub-configurations: an upright orientation, in which the globular density extends away from the lattice, and a bent orientation, in which the density is directed toward the MT surface (Figure 4D,E,F). Together, these data indicate that full-length KIF21B can adopt multiple MT-bound configurations compatible with both single-MT engagement and inter-MT crosslinking.

**Figure 4.**
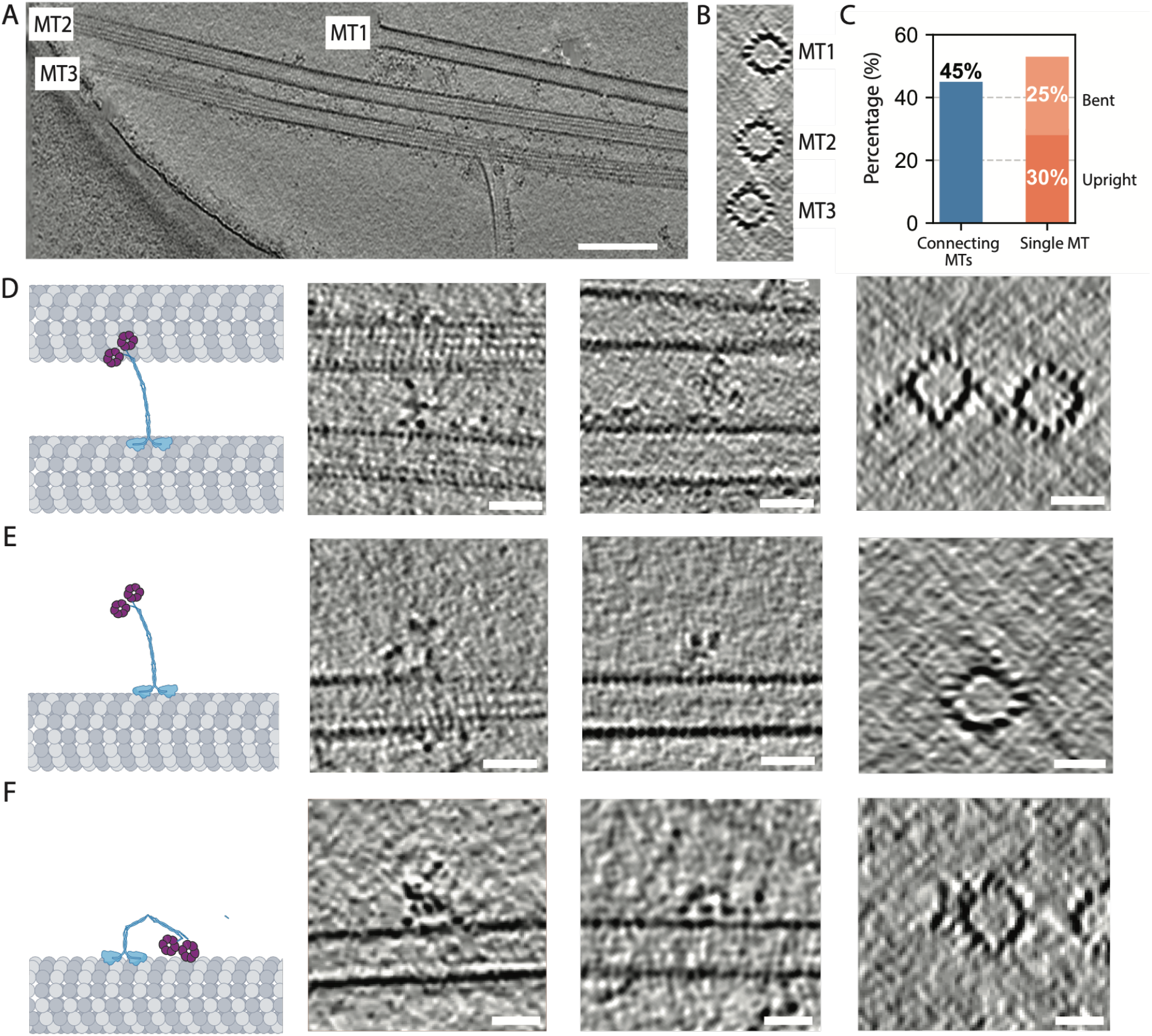
MT binding configurations of full-length KIF21B. (A) Section of a tomogram of GMPCPP MTs incubated with purified full-length KIF21B. Scale bar, 100 nm. (B) Orthogonal slice sums confirming 14 protofilaments. (C) Histogram of orientation distribution of full-length KIF21B particles in the cryo-ET dataset (n = 50). (D) Representative tomographic slices showing KIF21B densities spanning two neighboring MTs, with cartoon depicting crosslinking. (E) Representative tomographic slices showing KIF21B in an upright configuration on a single MT, with cartoon illustration. (F) Representative tomographic slices showing KIF21B in a bent configuration on a single MT, with cartoon. Scale bars in D,E,F, 20 nm.

## DISCUSSION

Kinesin-4 family members KIF21A and KIF21B are among a small number of kinesins known to use their C-terminal tails for both MT dynamics regulation and MT crosslinking (Figure 5A). Unlike the largely unstructured or coiled-coil tails found in most other kinesins (Seeger et al., 2012), the KIF21A/B tails terminate in a WD40 β-propeller domain. Here we show that this β-propeller, together with a short N-terminal linker, engages the MT lattice through a longitudinal binding mode that simultaneously spans the intradimer and interdimer tubulin interfaces. This extended binding footprint provides a structural basis for previous deletion experiments showing that both the linker and the β-propeller contribute to MT binding, and for how the KIF21B tail can detect nucleotide dependent changes in the MT lattice (van Riel et al., 2017).

**Figure 5.**
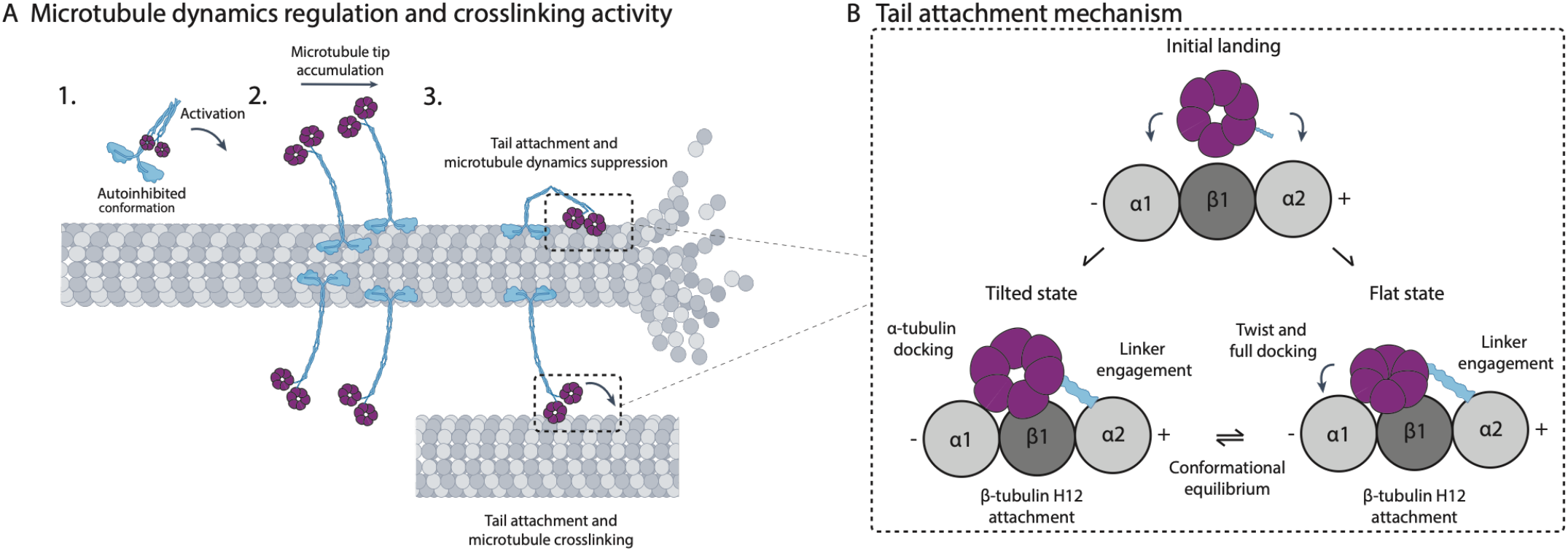
Model for tail-mediated KIF21B regulation of MT dynamics and MT crosslinking. (A) Proposed mechanism for MT binding and crosslinking by KIF21B. 1. KIF21B transitions from an autoinhibited to an active state (Cheng et al., 2014). 2. Activated KIF21B accumulates at the MT tip through plus-end directed movement. 3. Tail engagement with the lattice promotes stable tip association and MT dynamics regulation. Additionally, tail attachment to a second MT enables crosslinking. (B) Schematic model for tail attachment. The N-terminal linker engages the MT at an intradimer interface. Upon initial landing, the β-propeller can occupy a flat state in which initial contacts are established with α-tubulin and β-tubulin. The β-propeller can also occupy a tilted state, in which it is rotated ~35° relative to the β-tubulin surface.

The GTP bound MT lattice adopts an expanded lattice relative to the GDP state (Alushin et al., 2014). By bridging two longitudinally spaced tubulin binding sites, the linker at the intradimer interface and the β-propeller at the neighboring interdimer interface, the KIF21B tail is well positioned to detect nucleotide dependent lattice conformational differences. The preference of KIF21B for the GMPCPP/GTP-state lattice may reflect complementarity between the tail”s binding footprint and the expanded protofilament spacing of this state. This model is reminiscent of GTP-cap recognition by end-binding (EB) proteins, which similarly recognize conformational features of the GTP lattice to accumulate at growing MT ends (Bieling et al., 2007; Maurer et al., 2012).

Our cryo-EM data identify two engagement states rather than a single static binding pose. The β-propeller can occupy a tilted state and a fully engaged flat state, while the linker remains anchored at the intradimer pocket in both conformations. We propose a sequential docking model in which the linker first engages the MT at a hydrophobic intradimer pocket, providing a positional anchor. In this model, the β-propeller samples a tilted orientation through electrostatic interactions with the β-tubulin H12 helix before reaching a flat, more extensively engaged configuration in which it rotates ~35° (Figure 5B). Such a two-state binding model could allow the tail to sample the MT surface before committing to full engagement, a feature potentially important for tracking growing MT ends under dynamic conditions.

The cryo-ET data from full-length KIF21B illustrate how the two MT-binding modules, the tail and the motor domain, may be arranged to support distinct MT-bound configurations (Figure 5A,B). Bridging densities of ~28 nm connecting adjacent MTs with orientational flexibility are compatible with KIF21B acting as a flexible MT crosslinker. Configurations in which full-length KIF21B lies along a single MT are compatible with both binding modules engaging the same filament simultaneously, potentially stabilizing a compact, lattice-associated state. Further work will be needed to define the complete range of full-length KIF21B MT-binding geometries.

The MT contacting residues identified here are highly conserved between KIF21A and KIF21B and across species, suggesting that this tail attachment mechanism may be shared within the KIF21 subfamily (Supplementary Figure 3). A second MT-binding region within the coiled-coil 2 (CC2) segment of KIF21B has also been reported, raising the possibility that additional contacts cooperate with the distal tail to tune MT affinity, tip residence time, or crosslinking geometry (van Riel et al., 2017). Future structural studies of full-length KIF21A/B dimers in complex with MTs and regulatory partners such as KANK proteins will be important for understanding how distal tail–MT engagement is coordinated with motor activity, autoinhibition relief, and cortical anchoring (Weng et al., 2018; Guo et al., 2018; Haque et al., 2026). Such studies should help illuminate how KIF21A/B proteins shape dynamic MT arrays that underlie cell polarity, migration, axon guidance and synaptic function.

## ACKNOWLEDGEMENTS

We thank Dan Toso and the Cal-cryo facility at UC Berkeley for assistance with cryo-EM and cryo-ET data collection. We thank Ahmet Yilidz for feedback on figure designs. AR is thankful for the DBT Ramalingaswami Re-entry Fellowship.

## AUTHOR CONTRIBUTIONS

Conceptualization (A.T, B,L, J.P, A.A, E.N); Investigation (A.T, B.L, J.P, A.R, A.A, E.N); Writing – original draft (A.T, J.P, A.A, E.N).

## METHODS

### Protein Expression and Purification

The human KIF21BL–WD40–mNG construct (residues 1247–1674 with an N-terminal twin-Strep tag) was cloned into a pFastBac backbone. The plasmid was transformed into DH10Bac competent cells and plated on Bacmid selection plates with BluoGal at 37°C for three days. A white colony was selected, grown overnight in LB medium, and Bacmid DNA was isolated and transfected into adherent Sf9 cells using CellFectin. Transfected cells were incubated at 27°C for 5 days to produce the P1 virus. Two milliliters of P1 virus was added to 50 mL of suspension Sf9 culture and incubated for 5 days at 27°C to amplify the P2 virus, which was collected by centrifugation at 4,000 × g for 10 min and stored at 4°C in the dark. For protein production, 600 mL of Sf9 cells at 2 × 10^6^ cells/mL were infected with P2 virus at a 1:100 (v/v) ratio and incubated for 72 h at 27°C. Cells were harvested by centrifugation at 4,000 × g, and the pellet was snap-frozen in liquid nitrogen and stored at −80°C.

For purification, the cell pellet was thawed and resuspended in lysis buffer (50 mM PIPES-KOH pH 8.0, 300 mM KCl, 1 mM MgCl_2_, 1 mM DTT, 10% glycerol) and lysed by dounce homogenization. The lysate was cleared by centrifugation at 55,000 × g for 45 min in a Ti70 rotor (Beckman Coulter) and applied to 1 mL of Strep-Tactin XT resin pre-equilibrated in lysis buffer. The resin was washed with 10 column volumes of lysis buffer, and protein was eluted with lysis buffer supplemented with 100 mM D-biotin. The eluate was concentrated using a 50 kDa MWCO centrifugal concentrator and injected onto a Superdex 200 Increase 10/300 GL column equilibrated in lysis buffer. Peak fractions were pooled, concentrated to 6 µM, snap-frozen in liquid nitrogen, and stored at −80°C.

Purification of FL KIF21B followed the protocol established previously (van Riel et al., 2017). GFP-TEV-Bio–tagged constructs were co-transfected with BirA into HEK293T cells. Cells were lysed in buffer containing 50 mM HEPES, pH 7.4, 300 mM NaCl, 1 mM MgCl_2_, 0.5% Triton X-100, 1 mM DTT, and 1× cOmplete protease inhibitor cocktail (Roche). Lysates were incubated with M-280 Streptavidin Dynabeads (Invitrogen) for 1 h. Beads were washed three times with lysis buffer lacking protease inhibitor and then three additional times with cleavage buffer consisting of 50 mM HEPES, pH 7.4, 150 mM NaCl, 1 mM MgCl_2_, 0.05% Triton X-100, 1 mM DTT, and 1 mM EGTA. Bound proteins were released by incubation in cleavage buffer supplemented with 40 ng/µL TEV protease (770 nM; Sigma-Aldrich, St. Louis, MO) for 2 h at 4°C. The supernatant was collected and stored at −80°C until further use. Sample purity was assessed by SDS-PAGE followed by Coomassie staining. Stock concentrations of full-length KIF21B-GFP ranged from approximately 50 to 250 nM.

### Sample Preparation for Light Microscopy

Coverslips were prepared using a standard PEG–biotin passivation protocol. Plain glass coverslips were sequentially sonicated for 10 min each in water, acetone, and water, then sonicated for 40 min in 1 M KOH. After extensive rinsing with ddH_2_O, coverslips were silanized with APTES in an acetate/methanol solution for 10 min and washed with methanol. For PEG-biotinylation, 30 µL of 25% biotin-PEG-succinimidyl valerate in NaHCO_3_ buffer (pH 7.4) was sandwiched between two coverslips and incubated at 4°C overnight. Coverslips were rinsed with water, air-dried, vacuum-sealed, and stored at −20°C. Flow chambers were assembled by placing double-sided tape between a PEG-coated coverslip and a glass slide.

### Light Microscopy

Light microscopy was performed as previously described (Cetin and Taheri et al., 2025) on a custom-built multicolor, objective-type TIRF microscope based on a Nikon Ti-E body equipped with a 100× 1.49 NA apochromatic oil-immersion objective and the Nikon Perfect Focus System. Fluorescence was collected on an EMCCD camera (Andor iXon EM+, 512 × 512 pixels; effective pixel size 160 nm in the sample plane).

Alexa 488/GFP/mNeonGreen, LD555, and LD655 fluorophores were excited with 488, 561, and 633 nm lasers (Coherent) delivered through a single-mode fiber (Oz Optics), and emission was selected with 525/40, 585/40, and 697/75 bandpass filters (Semrock). Image acquisition was controlled with Micro-Manager v1.4. Flow chambers were incubated with 5 mg/mL streptavidin for 3 min, rinsed with BRB80 (80 Mm PIPES pH 6.8, 1 mM EGTA, 2 mM MgCl_2_), loaded with biotinylated MTs for 3 min, and washed with BRB80 supplemented with 1 mg/mL casein and 0.5% Pluronic acid. KIF21BL–WD40–mNG was diluted into the same buffer before introduction into the flow chamber.

### Fluorescence Image Analysis

Fluorescence images were imported into ImageJ and average MT fluorescence intensity was measured using the Measure tool. Data were plotted in Prism and fit to Y = B_max_× X / (Kdapp + X) to determine the apparent dissociation constant.

### Cryo-EM Sample Preparation

Lyophilized porcine brain tubulin (Cytoskeleton) was resuspended to 10 mg/mL in BRB80 supplemented with 10% glycerol, 1 mM GTP, and 1 mM DTT. Ten microliters of tubulin solution and 1 µL of 10 mM GMPCPP were polymerized at 37°C for ~2 h. MTs were pelleted at 37°C and 17,000 × g for 20 min, the supernatant discarded, and the pellet resuspended in BRB80 and diluted to 2.5 µM. All MT-binding proteins were desalted into BRB80 immediately before use using Zeba Spin columns (Pierce). For grid preparation, 2.5 µL of 2.5 µM GMPCPP-MTs were applied to a glow-discharged holey carbon grid (Quantifoil Cu 300 R1.2/1.3) for 30 s, manually blotted with Whatman filter paper, and 2.5 µL of 5 µM KIF21BL–WD40–mNG was added. Grids were transferred to a Vitrobot (Thermo Fisher Scientific) set at 22°C and 100% humidity and plunge-frozen in liquid ethane (blot force 5, blot time 6 s).

### Cryo-EM Data Collection

Cryo-EM data were collected on a Titan Krios microscope (Thermo Fisher Scientific) operated at 300 kV with a K3 direct electron detector (Gatan). Images were acquired at 105,000x magnification at 1.149 Å/pixel, with a dose rate of ~7.2 e^−^/pixel/s and an exposure time of ~9 s, dose-fractionated into 50 frames. Data collection used SerialEM (Mastronarde, 2003).

### Cryo-ET Sample Preparation

GMPCPP microtubules were prepared as described above. To prepare FL-KIF21B samples on microtubules for cryo-ET, in a 37°C chamber 2.5 µL of 20 µM GMPCPP-MTs were applied to a glow-discharged holey carbon cryo-EM grid (QuantiFoil, AU 200 R 2/2) and 0.5 ul of 20 uM unpolymerized tubulin in the presence of GTP was added and incubated for 30 s. Grids were subsequently manually blotted with Whatman filter paper, and 3 µL of 3 µM FL-KIF21B in the presence of 2mM AMPPNP were added to the grid, blotted again and another 3 ul of 3 µM FL-KIF21B with 2 mM AMPPNP were added to the grid.

During the last application, 2 ul of 5 times concentrated 10 nm BSA gold fiducials in BRB80 buffer were added onto the grid as well. Grids were then transferred to a Leica GP2 plunger preset to 37°C and blotted for 4.5 sec from the backside of the grid before immediately plunging into cooled liquid ethane.

### Cryo-ET Data Collection

Tilt series of FL-KIF21b bound to GMPCPP microtubules were collected on a Titan Krios microscope (Thermo Fisher Scientific), operated at 300 kV with a K3 direct electron detector (Gatan). Images were acquired at 33,000x nominal magnification at 2.63 A°/pixel and a defocus range of – 2 to – 5 µm. All data were collected using the SerialEM software package version 4.0 beta and the tilt series controller. Tilt series were collected as dose-symmetric from −60° to 60° using a 3° increment and total dose of 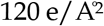 distributed equally across the tilt series and 5 frames per tilt.

### Cryo-ET Data Processing

Tilt series movies were motion corrected using Motioncor2 (Zheng et al., 2017) and tomograms were reconstructed in etomo in IMOD (Kremer et al., 1996). Within IMOD, tilt series alignment was performed by semi-automated gold fiducial tracking which was manually adjusted if necessary. For visualization, tomograms were binned two times and 3D Gaussian filtered. Tomograms were visualized in 3dmod IMOD using the slicer functionality to visualize tomograms along the direction of identified microtubules. Distance measurements were performed manually in IMOD. To analyze the distribution of different possible conformations on microtubules a total of 50 protein densities were visually classified either as connecting two neighboring microtubules, standing up on a single microtubule or bending down on a single microtubule.

### Cryo-EM Image Processing

Data processing followed established protocols for MT cryo-EM (Cetin et al., 2025; Zhang and Nogales, 2015). Movie stacks were motion-corrected in cryoSPARC (Punjani et al., 2017), and CTF parameters were estimated with the patch CTF job; micrographs with poor CTF fits were excluded manually. Particles were picked automatically with the filament tracer, initially without a template and subsequently using 2D class averages, with a segment separation of 82 Å. Particle images were extracted at 512 px and subjected to four rounds of 2D classification. Classes showing clear MT density were retained.

Heterogeneous refinement against 13- and 14-protofilament MT references was used to separate the two populations; the 14-protofilament class was used for all downstream processing. 14-protofilament particles were subjected to helical refinement (initial rise 82.5 Å, twist 0°) followed by local refinement with a cylindrical MT mask. Helical parameters were refined with the symmetry search job and used for pseudo-helical symmetry expansion. Local refinement on the expanded particle stack yielded the symmetrized MT reconstruction. The seam was identified without additional processing. Focused 3D classification on the KIF21B region above the MT surface resolved the tilted and flat engagement states, and independent local refinements yielded final maps at approximately 3.1 Å resolution.

### Model Building and Refinement

The KIF21B_L_–_W__D40_–_mNG_–MT structure was modeled by fitting the deposited GMPCPP-stabilized MT model (PDB: 6DPU; Zhang et al., 2018) and morphing with ISOLDE (Croll, 2018). The KIF21BL–WD40 region was predicted by AlphaFold3 (Jumper et al., 2021), fitted into the cryo-EM density, and refined with ISOLDE. The combined model was subjected to real-space refinement in PHENIX (Adams et al., 2010).

### Cryo-EM structure and data collection statistics

|  | KIF21B tail (flat) | KIF21B tail (tilted) |
| --- | --- | --- |
| EMDB ID | XXXXX | XXXXX |
| PDB ID | XXXX | XXXX |
| Data collection |  |  |
| Microscope | Titan Krios | Titan Krios |
| Camera | K3 | K3 |
| Voltage (kV) | 300 | 300 |
| Electron exposure (e<sup>-</sup>/Å<sup>2</sup>) | 50 | 50 |
| Magnification | 105,000x | 105,000x |
| Pixel size (Å/px) | 1.149 | 1.149 |
| Defocus range (μm) | -0.8 to -2.0 | -0.8 to -2.0 |
| Total micrographs | 814 | 814 |
| Final particles (post sym. expansion) | 299,514 | 264,926 |
| Map resolution (Å, FSC 0.143) | 3.05 | 3.06 |
| Sharpening B-factor (Å<sup>2</sup>) | 55 | 55 |
| Model refinement |  |  |
| Software | PHENIX | PHENIX |
| Non-hydrogen atoms | 12,720 | 12,608 |
| Protein residues | 1625 | 1608 |
| Ligands | 6 | 6 |
| Validation |  |  |
| Clashscore | 5.67 | 7.74 |
| MolProbity score | 1.71 | 1.95 |
| Rotamer outliers (%) | 1.9 | 2.21 |
| Ramachandran favored (%) | 96.89 | 96.10 |
| Ramachandran allowed (%) | 3.11 | 3.90 |
| Ramachandran outliers (%) | 0 | 0 |

**Supplementary Figure 1.**
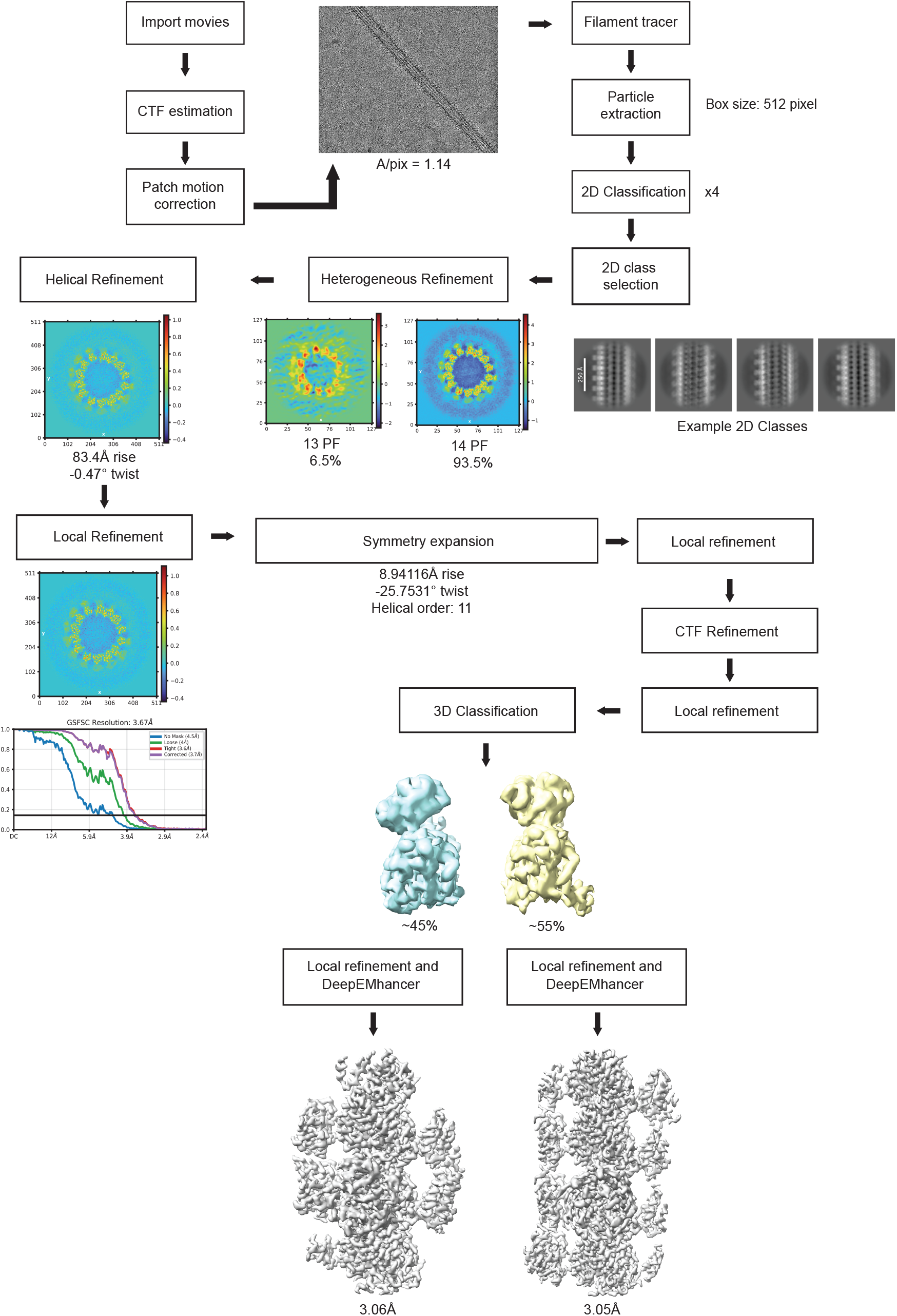
Cryo-EM Processing Pipeline. Overview of cryo-EM processing leading to the final reconstructions of KIF21B-bound MTs.

**Supplementary Figure 2.**
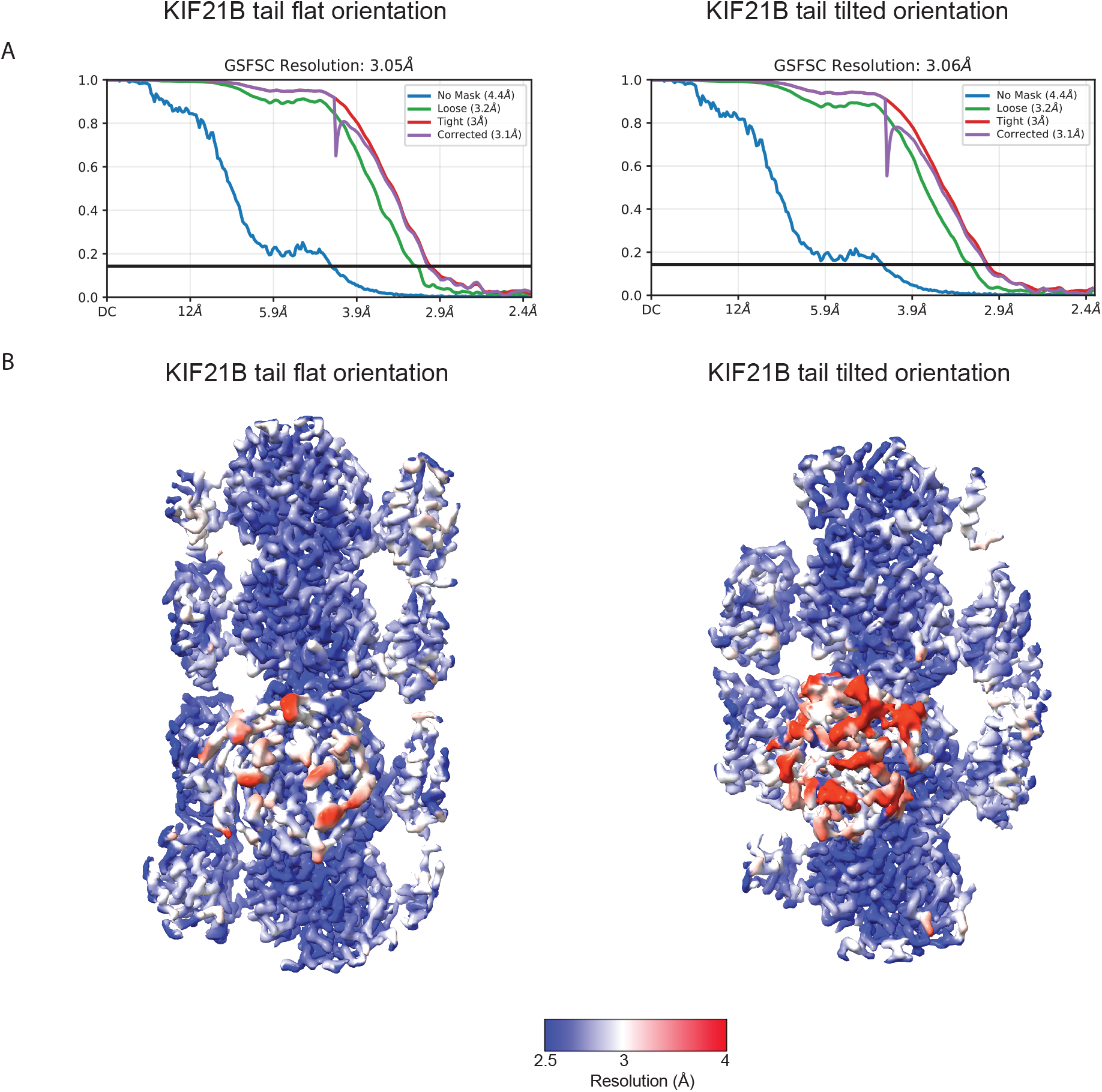
Local resolution mapping and FSCs. (A) FSC plots of the two MT bound KIF21B structures. (B) Local resolution maps of the two MT bound KIF21B structures

**Supplementary Figure 3.**
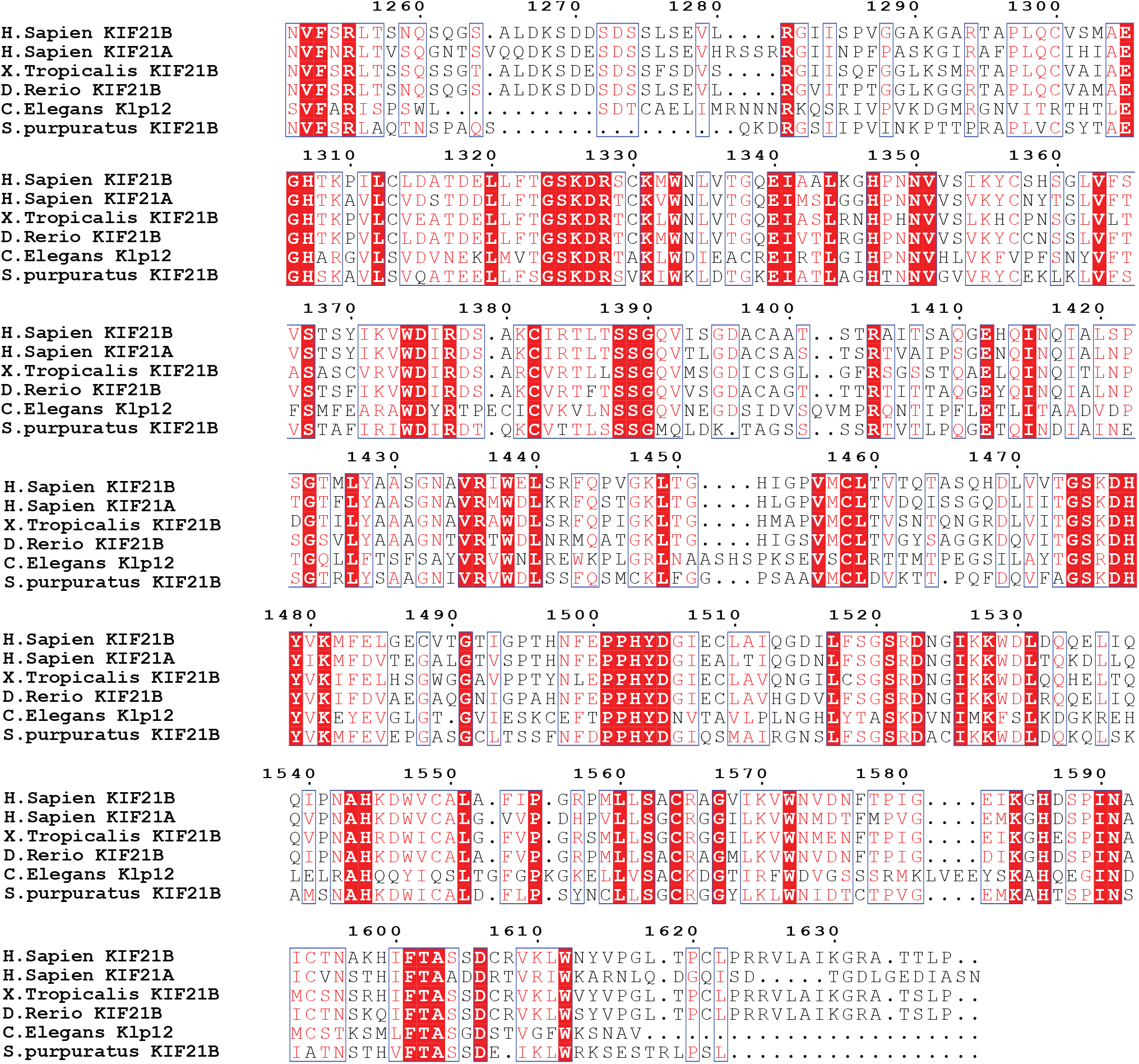
Sequence alignment of the KIF21B tail. Sequence alignment of the KIF21B and KIF21A tail across species. Alignment was displayed using ESPript3 (Robert et al., 2025).

## REFERENCES

Adams PD et al. (2010). PHENIX: a comprehensive Python-based system for macromolecular structure solution. Acta Crystallogr D Biol Crystallogr 66, 213–221. doi:10.1107/S0907444909052925.

Al-Haddad C, Boustany RM, Rachid E, Ismail K, Barry B, Chan WM, and Engle E (2021). KIF21A pathogenic variants cause congenital fibrosis of extraocular muscles type 3. Ophthalmic Genet 42, 195–199. doi:10.1080/13816810.2020.1852576.

Alushin GM, Lander GC, Kellogg EH, Zhang R, Baker D, and Nogales E (2014). High-resolution microtubule structures reveal the structural transitions in αβ-tubulin upon GTP hydrolysis. Cell 157, 1117–1129. doi:10.1016/j.cell.2014.03.053.

Asselin L et al. (2020). Mutations in the KIF21B kinesin gene cause neurodevelopmental disorders through imbalanced canonical motor activity. Nat Commun 11, 2441. doi:10.1038/s41467-020-16294-6.

Bianchi S et al. (2016). Structural basis for misregulation of kinesin KIF21A autoinhibition by CFEOM1 disease mutations. Sci Rep 6, 30668. doi:10.1038/srep30668.

Bieling P, Telley IA, and Surrey T (2010). A minimal midzone protein module controls formation and length of antiparallel microtubule overlaps. Cell 142, 420–432. doi:10.1016/j.cell.2010.06.033.

Bringmann H, Skiniotis G, Spilker A, Kandels-Lewis S, Vernos I, and Surrey T (2004). A kinesin-like motor inhibits microtubule dynamic instability. Science 303, 1519–1522. doi:10.1126/science.1094838.

Cetin B, Taheri A, Golcuk M, Monroy BY, Fernandes J, Ori-McKenney KM, Gur M, Nogales E, and Yildiz A (2025). Structure and mechanism of microtubule stabilization and motor regulation by MAP9. bioRxiv. doi:10.1101/2025.11.17.688911.

Cheng L et al. (2014). Human CFEOM1 mutations attenuate KIF21A autoinhibition and cause oculomotor axon stalling. Neuron 82, 334–349. doi:10.1016/j.neuron.2014.02.038.

Coy DL, Hancock WO, Wagenbach M, and Howard J (1999). Kinesin’s tail domain is an inhibitory regulator of the motor domain. Nat Cell Biol 1, 288–292. doi:10.1038/13001.

Croll TI (2018). ISOLDE: a physically realistic environment for model building into low-resolution electron-density maps. Acta Crystallogr D Struct Biol 74, 519–530. doi:10.1107/S2059798318002425.

Ghiretti AE, Thies E, Tokito MK, Lin T, Ostap EM, Kneussel M, and Holzbaur ELF (2016). Activity-dependent regulation of distinct transport and cytoskeletal remodeling functions of the dendritic kinesin KIF21B. Neuron 92, 857–872. doi:10.1016/j.neuron.2016.10.003.

Guo Q, Liao S, Zhu Z, Li Y, Li F, and Xu C (2018). Structural basis for the recognition of kinesin family member 21A (KIF21A) by the ankyrin domains of KANK1 and KANK2 proteins. J Biol Chem 293, 557–566. doi:10.1074/jbc.M117.817494.

Haque F, Srinivasu BY, Engen JR, Wales TE, and Subramanian R (2026). How the non-motile kinesin KIF7 adapts conserved kinesin principles for its function in Hedgehog signaling. bioRxiv, Version 1. doi:10.64898/2026.02.06.704517.

Hooikaas PJ, Damstra HGJ, Gros OJ, van Riel WE, Martin M, Smits YTH, van Loosdregt J, Kapitein LC, Berger F, and Akhmanova A (2020). Kinesin-4 KIF21B limits microtubule growth to allow rapid centrosome polarization in T cells. eLife 9, e62876. doi:10.7554/eLife.62876.

Jumper J et al. (2021). Highly accurate protein structure prediction with AlphaFold. Nature 596, 583–589. doi:10.1038/s41586-021-03819-2.

Kreft KL, van Meurs M, Wierenga-Wolf AF, Melief MJ, van Strien ME, Hol EM, Oostra BA, Laman JD, and Hintzen RQ (2014). Abundant KIF21B is associated with accelerated progression in neurodegenerative diseases. Acta Neuropathol Commun 2, 144. doi:10.1186/s40478-014-0144-4.

Kremer JR, Mastronarde DN, McIntosh JR (1996). Computer visualization of three-dimensional image data using IMOD. J Struct Biol 116, 71–76. doi:10.1006/jsbi.1996.0013.

Marszalek JR, Weiner JA, Farlow SJ, Chun J, and Goldstein LSB (1999). Novel dendritic kinesin sorting identified by different process targeting of two related kinesins: KIF21A and KIF21B. J Cell Biol 145, 469–479. doi:10.1083/jcb.145.3.469.

Mastronarde DN (2003). SerialEM: a program for automated tilt series acquisition on Tecnai microscopes using prediction of specimen position. Microsc Microanal 9, 1182–1183. doi:10.1017/S1431927603445911.

Maurer SP, Fourniol FJ, Bohner G, Moores CA, Surrey T (2012). EBs recognize a nucleotide-dependent structural cap at growing microtubule ends. Cell 149, 371–382. doi:10.1016/j.cell.2012.02.049.

Mazumdar M, Sundareshan S, and Misteli T (2004). Human chromokinesin KIF4A functions in chromosome condensation and segregation. J Cell Biol 166, 613–620. doi:10.1083/jcb.200401142.

Muhia M et al. (2016). The kinesin KIF21B regulates microtubule dynamics and is essential for neuronal morphology, synapse function, and learning and memory. Cell Rep 15, 968–977. doi:10.1016/j.celrep.2016.03.086.

Nithianantham S et al. (2023). The kinesin-5 tail and bipolar minifilament domains are the origin of its microtubule crosslinking and sliding activity. Mol Biol Cell 34, ar111. doi:10.1091/mbc.E23-07-0287.

Park H et al. (2025). The kinesin-4 protein KIF27 forms a cytoskeletal scaffold at the transition zone to promote motile cilia structural integrity. Proc Natl Acad Sci USA 122, e2515392122. doi:10.1073/pnas.2515392122.

Punjani A, Rubinstein JL, Fleet DJ, and Brubaker MA (2017). cryoSPARC: algorithms for rapid unsupervised cryo-EM structure determination. Nat Methods 14, 290–296. doi:10.1038/nmeth.4169.

Robert, X., Guillon, C. and Gouet, P. (2025) FoldScript: a web server for the efficient analysis of AI-generated 3D protein models. Nucleic Acids Res. 53(W1), W277–W282. doi: 10.1093/nar/gkaf326

Seeger MA, Zhang Y, and Rice SE (2012). Kinesin tail domains are intrinsically disordered. Proteins 80, 2437–2446. doi:10.1002/prot.24128.

Subramanian R, Ti SC, Tan L, Darst SA, and Kapoor TM (2013). Marking and measuring single microtubules by PRC1 and kinesin-4. Cell 154, 377–390. doi:10.1016/j.cell.2013.06.021.

Taguchi S et al. (2022). Structural model of microtubule dynamics inhibition by kinesin-4 from the crystal structure of KLP-12–tubulin complex. eLife 11, e77877. doi:10.7554/eLife.77877.

Vale RD, Schnapp BJ, Reese TS, and Sheetz MP (1985). Movement of organelles along filaments dissociated from the axoplasm of the squid giant axon. Cell 40, 449–454. doi:10.1016/0092-8674(85)90159-X.

van der Vaart B et al. (2013). CFEOM1-associated kinesin KIF21A is a cortical microtubule growth inhibitor. Dev Cell 27, 145–160. doi:10.1016/j.devcel.2013.09.010.

van Riel WE, Rai A, Bianchi S, Katrukha EA, Liu Q, Heck AJR, Hoogenraad CC, Steinmetz MO, Kapitein LC, and Akhmanova A (2017). Kinesin-4 KIF21B is a potent microtubule pausing factor. eLife 6, e24746. doi:10.7554/eLife.24746.

Weng Z, Shang Y, Yao D, Zhu J, and Zhang R (2018). Structural analyses of key features in the KANK1·KIF21A complex yield mechanistic insights into the cross-talk between microtubules and the cell cortex. J Biol Chem 293, 215–225. doi:10.1074/jbc.M117.816017.

Wickstead B and Gull K (2006). A “holistic” kinesin phylogeny reveals new kinesin families and predicts protein functions. Mol Biol Cell 17, 1734–1743. doi:10.1091/mbc.e05-11-1090.

Yildiz A (2025). Mechanism and regulation of kinesin motors. Nat Rev Mol Cell Biol 26, 86–103. doi:10.1038/s41580-024-00780-6.

Yount AL, Zong H, and Walczak CE (2015). Regulatory mechanisms that control mitotic kinesins. Exp Cell Res 334, 70–77. doi:10.1016/j.yexcr.2014.12.015.

Yue Y, Blasius TL, Zhang S, Jariwala S, Walker B, Grant BJ, Cochran JC, and Verhey KJ (2018). Altered chemomechanical coupling causes impaired motility of the kinesin-4 motors KIF27 and KIF7. J Cell Biol 217, 1319–1334. doi:10.1083/jcb.201708179.

Zhang R, LaFrance B, and Nogales E (2018). Separating the effects of nucleotide and EB binding on microtubule structure. Proc Natl Acad Sci USA 115, E6191–E6200. doi:10.1073/pnas.1802637115.

Zhang R and Nogales E (2015). A new protocol to accurately determine microtubule lattice seam location. J Struct Biol 192, 245–254. doi:10.1016/j.jsb.2015.09.015.

Zheng SQ, Palovcak E, Armache JP, Verba KA, Cheng Y, Agard DA (2017). MotionCor2: anisotropic correction of beam-induced motion for improved cryo-electron microscopy. Nat Methods 14, 331–332. doi:10.1038/nmeth.4193.

