## Supplement for "Microtubule Engagement by the WD40-Containing Tail of Kinesin-4 KIF21B"

### Supplementary Figures

Aryan Taheri[1], Benjamin LaFrance[1,2], Julia Peukes[1,3], Ankit Rai[4,5], Anna Akhmanova[4], and Eva Nogales[1,6,7*]

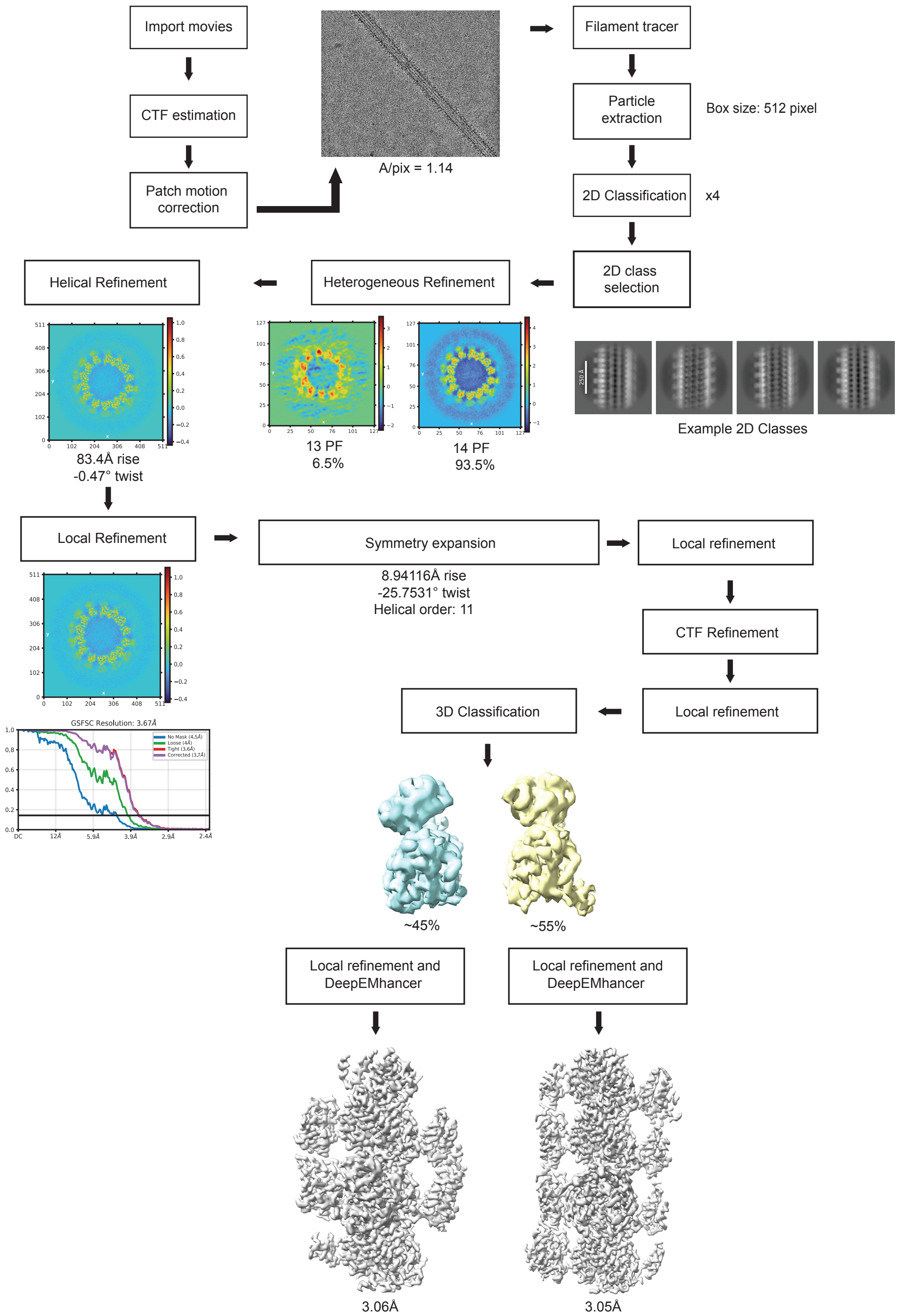


**Supplementary Figure 1.** | **Cryo-EM Processing Pipeline.** Overview of cryo-EM processing leading to the final reconstructions of KIF21B-bound MTs.

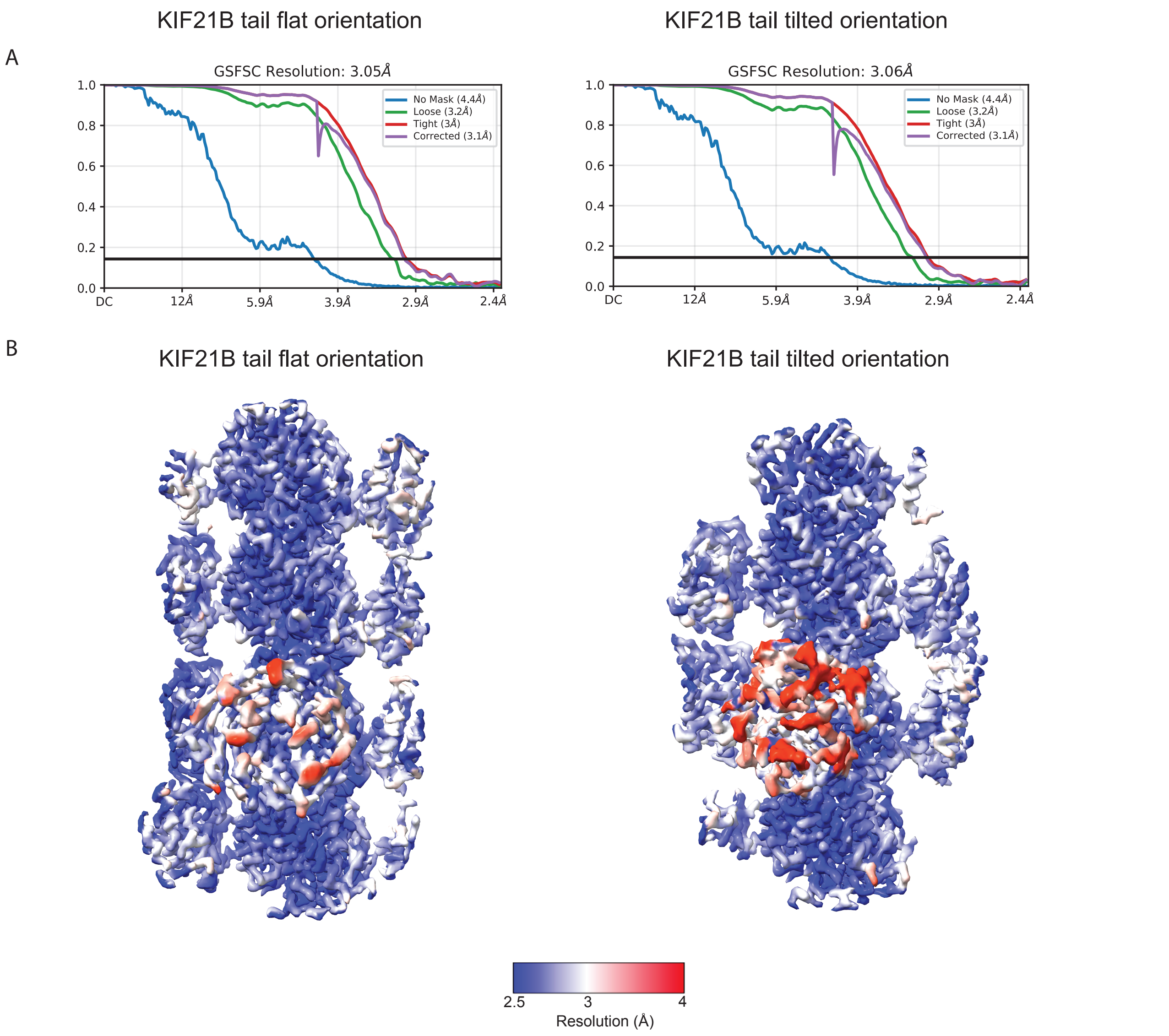


**Supplementary Figure 2.** | **Local resolution mapping and FSCs.** (A) FSC plots of the two MT bound KIF21B structures. (B) Local resolution maps of the two MT bound KIF21B structures

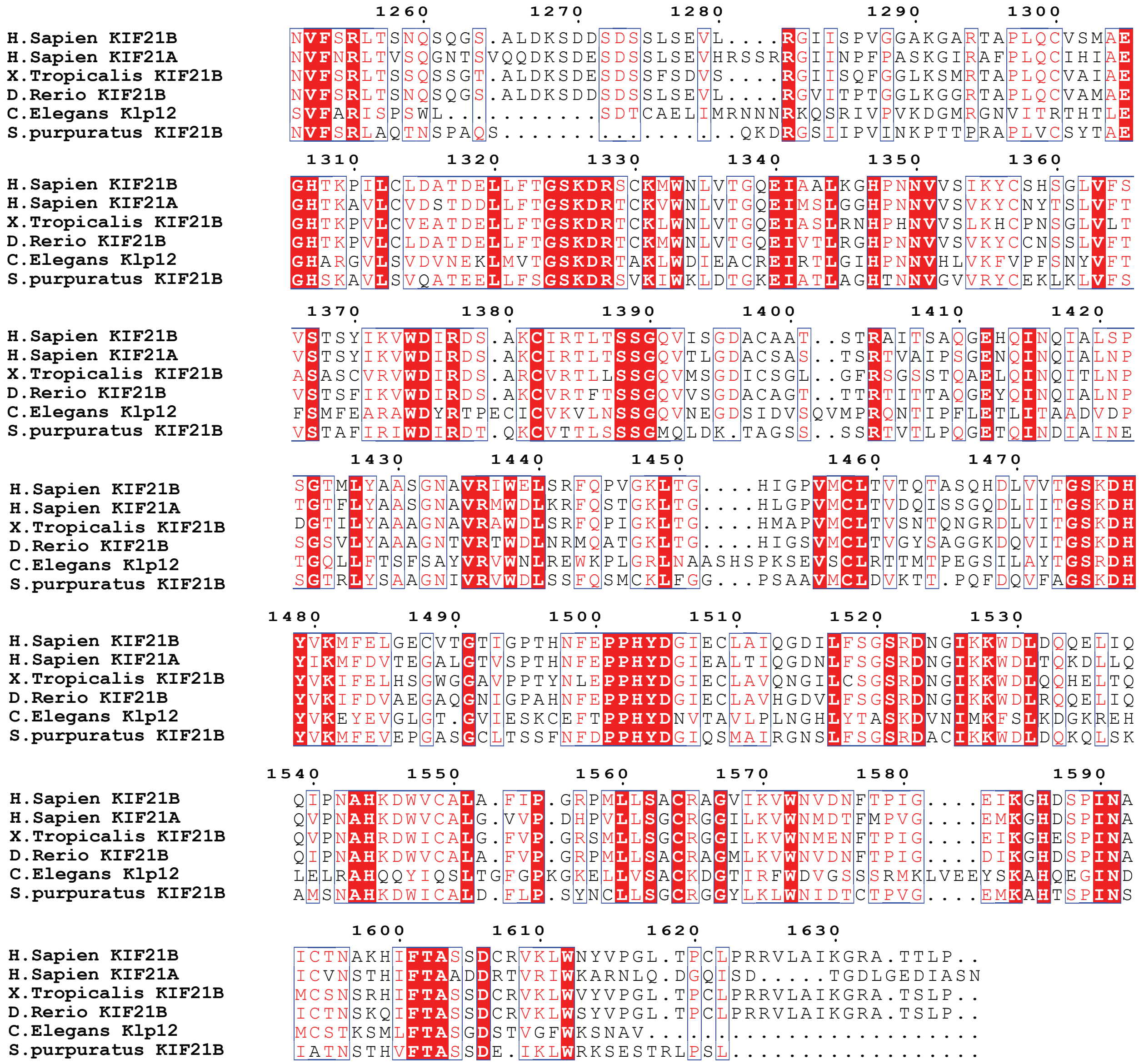


**Supplementary Figure 3. | Sequence alignment of the KIF21B tail.** Sequence alignment of the KIF21B and KIF21A tail across species. Alignment was displayed using ESPript3 (Robert et al., 2025).
